# Identification of the relative timing of infectiousness and symptom onset for outbreak control

**DOI:** 10.1101/571547

**Authors:** Robert C. Cope, Joshua V. Ross

**Affiliations:** The University of Adelaide, Stochastic Modelling & Operations Research Group, School of Mathematical Sciences, Adelaide SA 5005, AUSTRALIA.

**Keywords:** Bayesian model discrimination, epidemiology, optimal experimental design, random forests

## Abstract

In an outbreak of an emerging disease the epidemiological characteristics of the pathogen may be largely unknown. A key determinant of ability to control the outbreak is the relative timing of infectiousness and symptom onset. We provide a method for identifying this relationship with high accuracy based on data from household-stratified symptom-onset data. Further, this can be achieved with observations taken on only a few specific days, chosen optimally, within each household. This constitutes an important tool for outbreak response. An accurate and computationally-efficient heuristic for determining the optimal surveillance scheme is introduced. This heuristic provides a novel approach to optimal design for Bayesian model discrimination.

## Introduction

The timing of infectiousness relative to symptom onset has been identified as a key factor in ability to control an outbreak (*11*). The explanation is intuitive: If symptoms appear before infectiousness, then contact tracing and isolation strategies will be effective, whereas for post-infectiousness symptom presentation, broader, non-symptom based strategies must be adopted. Consequently, identifying the relative timing as early as possible in an outbreak is imperative to assessing potential for control and selecting a measured response.

Severe acute respiratory syndrome (SARS) is a prime example of a disease in which symptoms foreshadow significant levels of infectiousness (*2*). This played a critical role in limiting mortality and morbidity in outbreaks during 2003, via simple public health measures such as isolation and quarantining (*2,8,11,14,16,18*). Smallpox is most similar to SARS in this respect, but must be contrasted with HIV, where a large proportion of secondary infections occur before symptoms (*11*). For influenza, the relationship is less clear, with symptoms and infectiousness likely coinciding closely, with some transmission possible before symptom onset (*17, 21,*). This relationship will not be known in an outbreak of an emerging pathogen, and one must turn to early outbreak surveillance data for insights.

Many jurisdictions organize their emerging disease monitoring policies around households. As an example, First Few Hundred studies are proposed as a first response surveillance scheme following the identification of a novel disease and/or strain as part of national pandemic plans (*3, 12, 19*). Following the observation of a first symptomatic individual, their household is enrolled in an intensive surveillance program, so that day of symptom onset for subsequent cases within that household are recorded. Methods have recently been developed to characterise transmissibility and severity of a novel pathogen – other factors influencing ability to control an outbreak (*11*) – based on such data. Currently lacking is a method for accurate determination of relative timing of infectiousness and symptom onset using this data.

Here we introduce, and demonstrate through a simulation study, a method for identifying with high accuracy the timing of infectiousness relative to symptom onset from household-stratified symptom surveillance data. Remarkably, we show this is achievable with observations taken on only a few specific days, chosen optimally, within each household. Our approach to determining the optimal surveillance scheme is based on an efficient heuristic. This heuristic provides a general, computationally-efficient approach to optimal design for Bayesian model discrimination.

## Bayesian model discrimination for outbreak control

We model disease dynamics within each household as a continuous-time Markov chain (*15*), that counts the number of household members that are susceptible (S), exposed (E), infectious (I), or recovered (and immune; R). Under this model, the timing of symptom onset relative to infectiousness is mapped to which transition is observed: symptoms appear either upon infection, infectiousness, or recovery. The challenge is to determine which of these three (observation) models best describes the household-stratified symptom-onset data (Figure 1a).

There is a relatively rich literature on Bayesian model discrimination (*1, 7, 10, 29*), and optimal design for such (*6, 28,*), which are the most appropriate tools and framework to address this question. A general difficulty with this theory is that practical implementation is at best difficult, and often infeasible. This has led to methods based on approximate Bayesian computation (ABC), which requires only simulation of realisations from each model, and is computationally feasible for a wide range of models. Unfortunately, there exists ‘a fundamental difficulty’ in establishing robust methods based upon summary statistics (*25, 26,*); however, see the recent work of Dehideniya *et al.* (*9*).

Another approach to model discrimination in an ABC framework has been proposed by Pudlo *et al.* (*22*). They treat model discrimination as a classification problem, for which machine learning methods are ideal, and in particular propose the use of random forests to perform this task. This approach provides a highly-efficient, and importantly, robust method for model discrimination. Hainy *et al.* (*13*) expand on this approach as specifically applied to optimal design for model discrimination.

We apply these tools, first for accurate, robust characterisation of relative timing of symptoms and infectiousness, and second, for optimal design of early outbreak surveillance for accurate model discrimination. Specifically, the aim of the latter is to select an optimal surveillance scheme, consisting of a fixed number of observations, in order to discriminate three different timings of symptom onset relative to infectiousness, within a household-stratified epidemic model. We evaluate the impact of assumptions and summary statistics. Additionally, we propose a new, computationally-efficient and highly-accurate heuristic for optimal design choice, which in this application determines the optimal days upon which to perform surveillance in households.

## Methods

### Epidemic model

We demonstrate using an example system of a novel infectious disease, spreading in a population structured into households. We assume that the population is large and mixing between households is random, such that after a household is initially infected, the remaining transmission within the household is independent of transmission outside the household (*5, 27*). Therefore, transmission dynamics within households can be modelled independently (*4*). Given this novel etiological agent, we wish to determine if symptom onset occurs at the time of infection, infectiousness, or recovery (i.e., these are the three candidate models we wish to discriminate). The model behaviours are otherwise assumed identical. To be emphatic, the underlying disease dynamics is identical in all three models, each differing only in when observations are made, corresponding to different timings of symptom onset (Figure 1a).

We model the epidemic dynamics in households as a continuous-time Markov chain (Figure 1a) (*15*). Individuals transition from susceptible (S) to exposed (E), then to infectious (I), and finally to recovered (R), with rates as described in Table 1.

**Table 1:** Events, transitions and rates within a household.

| Description | Transition | Rate |
| --- | --- | --- |
| Infection | $(S, I) \rightarrow (S - 1, E + 1)$ | $\beta SI / (N - 1)$ |
| Infectiousness | $(E, I) \rightarrow (E - 1, I + 1)$ | $\sigma E$ |
| Recovery | $(I, R) \rightarrow (I - 1, R + 1)$ | $\gamma I$ |

We assign a distribution to each parameter (Supplemental Figure S1), based on physical quantities to reflect the assumed prior knowledge of the etiological agent:

- 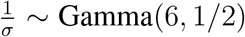 Gamma(6, 1/2), representing a mean exposed duration of 3 days (mode at approximately 2.5 days);
- 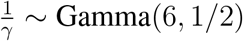 Gamma(6, 1/2), representing a mean infectious duration of 3 days (mode at approximately 2.5 days); and,
- *R*_0_ ~ 1 + Gamma(2, 1/2), representing a mean *R*_0_ (the expected number of secondary cases caused by an infectious individual in a fully susceptible population) of 2 (mode at approximately 1.5).

These distributions are sampled per-simulation, i.e., sampled parameters are kept constant across all households within a given epidemic. We also test the accuracy of model discrimination when these parameters are known, fixed quantities (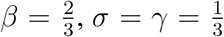; see Supplemental Figure S3 for results.

Following the first symptomatic case in a household, the number of symptomatic cases within the household is observed daily; i.e., the instant that the first individual in a household shows symptoms is time zero. Then, the number of cases seen before time 1 constitutes the first observation, between time 1 and 2 the next observation, and so on. This proceeds for 14 days, with any symptoms occurring after time 14 not observed.

When testing the effect of asymptomatic infections on model discrimination, we sample an additional parameter, *p*_obs_, the probability that an individual shows symptoms at the time they would in the model in question. We explored two scenarios: (1) *p*_obs_ ~ Beta(5, 5) (i.e., a mean *p*_obs_ of 0.5), and (2) *p*_obs_ ~ Beta(7.5, 2.5) (i.e., a mean *p*_obs_ of 0.75).

### Random forest model selection

To attempt to discriminate models, we use the approximate Bayesian random forest approach of Pudlo *et al.* (*22*). This proceeds as follows:

- Select a number of simulations, *N*_*s*_, and a number of households, *N*_*h*_.
- For each model:

– Sample a set of parameters *θ* = (*R*_0_, *σ*, *γ*) from the (prior) distributions.
– Simulate *N*_*h*_ households given these parameters.
– Repeat this process *N*_*s*_ times.
- Given the *N*_*s*_ simulations from each model, extract the data corresponding to the considered design.
- Construct a random forest that predicts the model label, given the simulations.
- Assess the accuracy of the process on a left-out test set.

Once a design has been chosen, to employ this process when an outbreak is observed it would be input to the (trained) random forest, to obtain a prediction of which model it is most consistent with.

Random forests were constructed using the Python scikit-learn RandomForestClassifier algorithm (*23*), with 200 trees.

### Summary statistics

To more effectively use the household data in training the random forest, we summarize raw household data as daily histograms of incidence, as in Figure 1c. That is, we count the proportion of households that, on day *d*, observed an incidence of *i*, and then use the resultant (design size) × (household size + 1) data vector as the new random forest predictors. For example, with designs of size 5, households of size 5, and 200 households, the raw data would consist of 5 × 200 = 1000 predictors, whereas the histogram summaries would consist of 5 × 6 = 30 predictors.

### Optimal sampling design

Conducting a First Few Hundred-style study can be extremely labour intensive. Consequently, we wish to assess the potential for model discrimination when sampling is only performed on a subset of days, rather than every day. If we choose to only sample on *D* < 14 days, within the first 14 days following the first symptomatic case in each household, we must necessarily also choose the optimal days on which to sample. We choose those days that produce the highest classification accuracy on a left-out test set. This design problem is small, with only 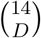 designs of size *D* (or 2^14^ = 16, 384 total designs) to evaluate, so we apply exhaustive search in this case; however a combinatorial optimisation algorithm could be applied and would likely be necessary in a more complex design problem to search for the optimal design.

### Heuristic solution

Rather than evaluating the full set of possible designs, or applying an optimisation algorithm, we propose a heuristic for efficiently finding high-quality designs of a given size. This heuristic is to perform random forest model selection on the largest possible design, extract the random forest feature importance Figure 1b), and use this random forest feature importance to rank design points. Specifically, days are ranked on their maximum feature importance; the sum of the importance of features from a day was also tested, but had inferior performance. A design of size *d* uses the highest-ranked *d* design points. The random forest feature importance metric we use is the mean decrease in Gini impurity (*24*) of a feature across the trees in the random forest (this metric is easily extracted from the python scikit-learn random forest algorithm (*23*)).

## Results

Random forest-based Bayesian model discrimination was able to accurately discriminate relative timing of symptoms and infectiousness for simulated household-stratified symptom-onset data: with 200 households of size 5, accuracy was 0.923 (with random parameters, and 10,000 training simulations per model). Accuracy was reduced with fewer households: to 0.853 with 100 households, and 0.657 with only 25 households (Figure 1d). These results were robust with respect to variation in household size, with accuracy ranging from 0.892 with 200 households of size 3 to 0.935 with 200 households of size 7.

Remarkably, model discrimination remained accurate when only a small subset of daily household data were observed, when the observations were from an optimal design: a design of size 5 and 190 households was sufficient to produce a classification accuracy of ≥ 0.90 (Figure 1d, Figure 2a). Accuracy increased as the design size (i.e., number of days of surveillance) and the number of households increased. The heuristic produced the exact optimal design at design sizes 4 and 5 (Figure 2b), and an effectively indistinguishable level of accuracy compared to the optimal for larger design sizes (Figure 1d). The heuristic ensured a substantial reduction in computation time: to produce Figure 1d, 39 random forests were required when using the heuristic, compared to 49,107 random forests to produce the optimal results. We also explored the impact of varying household size, the amount of training data used, and of using fixed, known parameters rather than parameters sampled from a distribution: larger households and more training points produced small increases in accuracy (Supplemental Figures S2, S3), and known epidemic parameters produced substantial increases in accuracy (Supplemental Figure S3).

The key design points (i.e., sampling days) for optimal designs were consistently the first day (Figure 2b), followed by other days early in the outbreak (i.e., days 2–4), and the final sampling day (day 14). Days 6–13 typically had little impact on model discrimination accuracy (i.e., optimal Accuracy consistently levelled off as design size increased beyond 5; Figure 1d, Supplemental Figure S3), and the optimal combination of these days varied due to stochasticity in both training and test data. This is consistent with the feature importance used to develop the heuristic (Figure 1b), i.e., those days that were consistently optimal were those with highest feature importance.

To assess the impact of asymptomatic infections on model discrimination, we repeated the analysis, except with each individual only being symptomatic (at the point that they otherwise would) with probability *p*_obs_ (again, sampled from a prior distribution). This partial observation made model discrimination substantially more challenging: with designs of size 5 and 200 households (Figure 2a), accuracy was 0.796 when *p*_obs_ had a mean of 0.75, and accuracy was 0.653 when *p*_obs_ had a mean of 0.5 (compared to 0.908 with complete observation).

## Discussion

Identifying the relative timing of symptom onset and infectiousness in an emerging epidemic is critical to outbreak control. We have demonstrated that it is not only possible to accurately identify the relative timing based upon household-stratified data available early in an outbreak, but that it can be done without observing each household every day. Moreover, we can use random forest feature importance to inform a heuristic that vastly reduces the computation necessary to choose high-accuracy designs.

It is remarkable that it is possible to discriminate models so accurately, given that they share identical epidemic dynamics, and only differ in observation. The non-parametric nature of the random forest is able to use small but clear differences between models (e.g., Figure 2c) to extract sufficient information to discriminate them. Combining the raw household data to form summary statistics is critical to this: if the raw household data is used rather than the summary statistics, accuracy is substantially lower (Supplemental Figure S4). While it can be difficult to interpret the classifications made by a random forest-classifier, interrogating key individual predictors (as in Figure 2c) provides clarity, and elucidates why feature importance provides a useful heuristic for choosing optimal designs (*20*).

**Figure 1:**
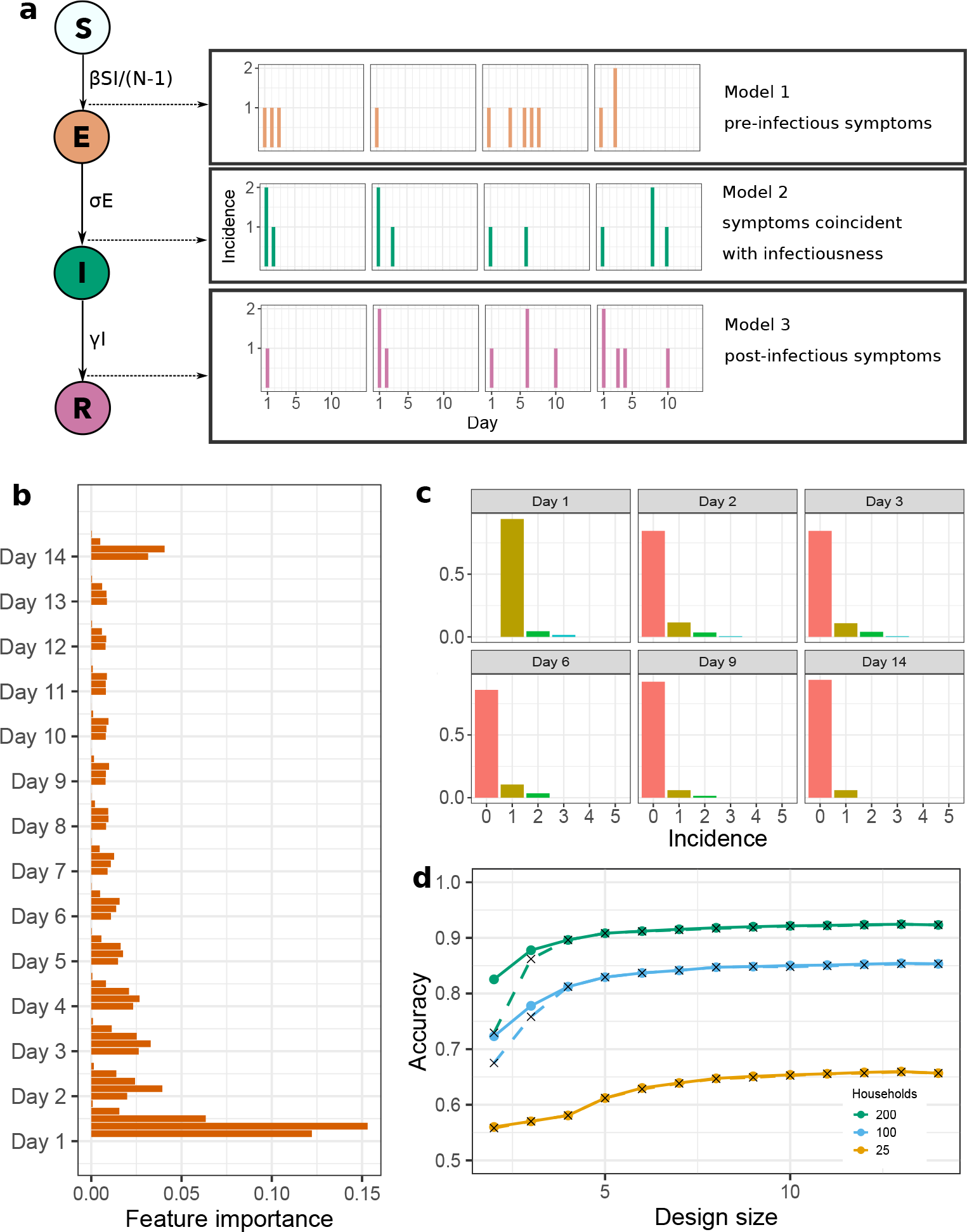
(a) Model schematic describing: transitions between states within each household continuous-time Markov chain; the three observation models being discriminated between; and, the way that these household-level data are observed. (b) Random forest feature importance for the full 14-day design, used to construct the heuristic for smaller designs. (c) Histogram summaries of the daily household-level data under a given design, used as predictors in the random forest. (d) Resulting random forest accuracy as design size increases, for the true optimal design (solid lines) and heuristic solution (crosses with dashed line). These results correspond to households of size 5, with 10,000 training samples from each model, each with parameters drawn from the distributions displayed in Supplemental Figure S1.

**Figure 2:**
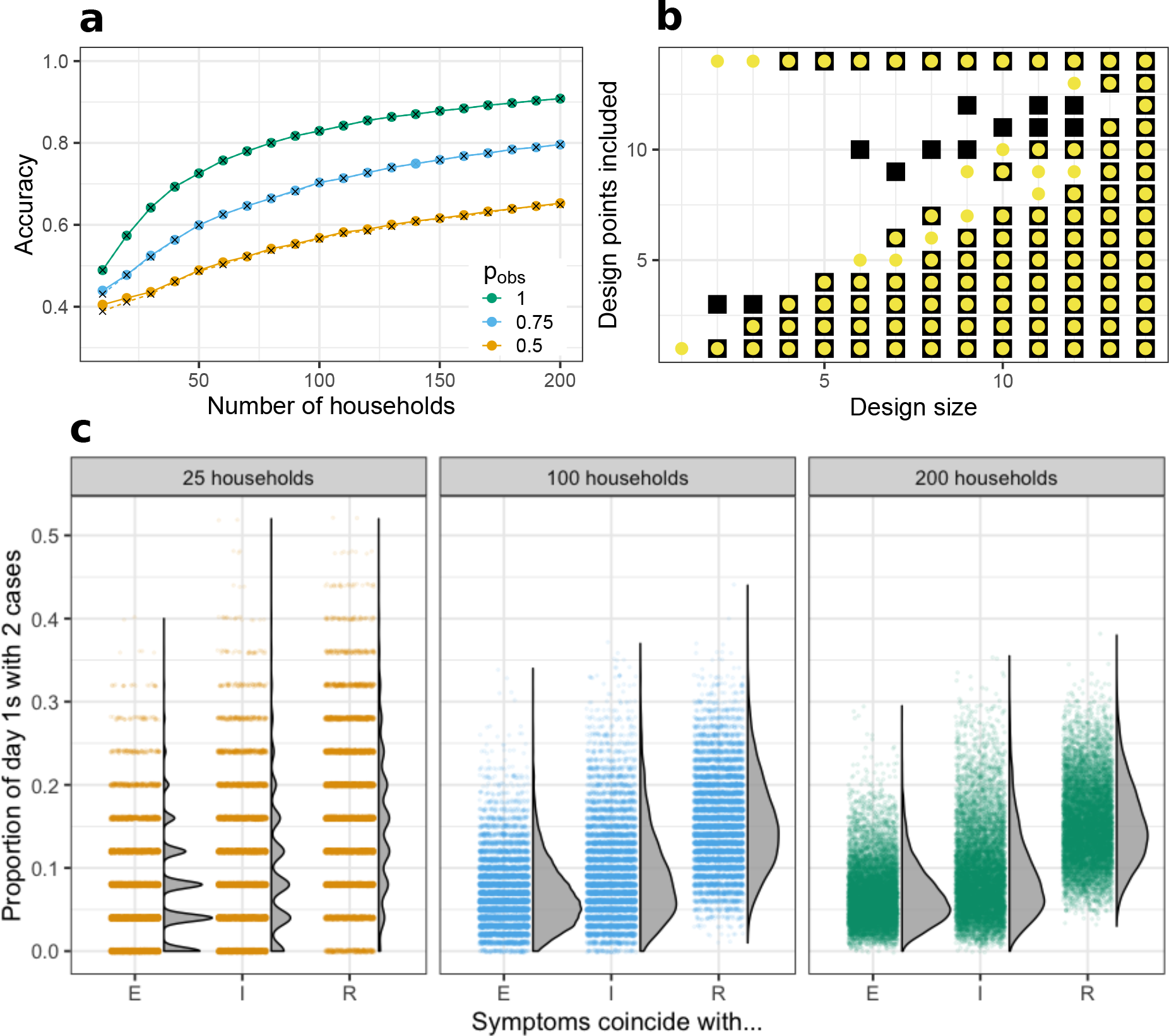
(a) Accuracy of model discrimination in designs of size 5, as the number of households increases, and under partial observation. Note that *p*_obs_ is not a fixed parameter but is sampled from a distribution; the listed value is its mean. The case with mean *p*_obs_ of 0.5 was sampled from a Beta(5,5) distribution, and a mean *p*_obs_ of 0.75 from a Beta(7.5,2.5) distribution. (b) Difference between heuristic designs (coloured points) and optimal designs (black boxes) as the design size increases. Note that the heuristic selects the optimal design at design sizes 4, 5, 13, and 14. (c) Distribution of training sample observations (under each model and number of households) for the most important feature under the heuristic: the proportion of households with 2 cases observed on day 1. These results correspond to households of size 5, with 10,000 training samples from each model, each with parameters drawn from the distributions that appear in Supplemental Figure S1.

The accuracy of model discrimination decreases substantially as the proportion of cases that are asymptomatic increases. However, this can be compensated by increasing the number of households. The outbreaks in which early control is most critical are likely to be those in which most individuals are symptomatic, due to symptoms being strongly correlated with severity, for example hospitalisations and deaths.

In some situations it may be necessary to consider more complicated surveillance schemes, in which case it may not be possible to evaluate the exact optimal design by exhaustive search. However, the heuristic proposed here should remain effective in more complicated design spaces, provided they have a similar form, i.e., designs of a given size are a subset of designs of larger sizes upon which the random forest can be trained to extract feature importance.

Assumptions impact any model-based study. Here assumptions include: a constant household size; enrolling each household in the study at the instant its first member shows symptoms; and, most critically, assuming that the underlying epidemic model is true. It is possible to select between models that differ in addition to observation process; however any increase in the number of models to classify will likely result in increased computation and potentially decreased accuracy.

In the future, the aim is to combine Bayesian model discrimination and parameter estimation in an online manner. Improving estimates of parameters improves the ability to discriminate models, and, more certainty regarding the model likely reduces variance in parameter estimates. This would allow for unified characterisation of all factors influencing the ability to control an outbreak.

## Supplemental Information

The supplemental information contains:

- Plots of the prior distribution for each epidemic parameter used to generate the household data (Fig. S1).
- A comparison of the accuracy of model discrimination as the size of households in the model varies from 3 to 7. This includes both the complete observation scenarios, and the scenarios wherein *p*_obs_ = 0.5 (Fig. S2).
- A comparison of the accuracy of model discrimination when parameters are known (fixed) values versus values sampled from the prior distributions; and of the impact of using 1,000 versus 10,000 training samples (Fig. S3).
- Model discrimination accuracy when the random forests are trained on the raw, unsummarised data rather than the histogram summaries that appear in the main text (Fig. S4).

**Figure S1:**
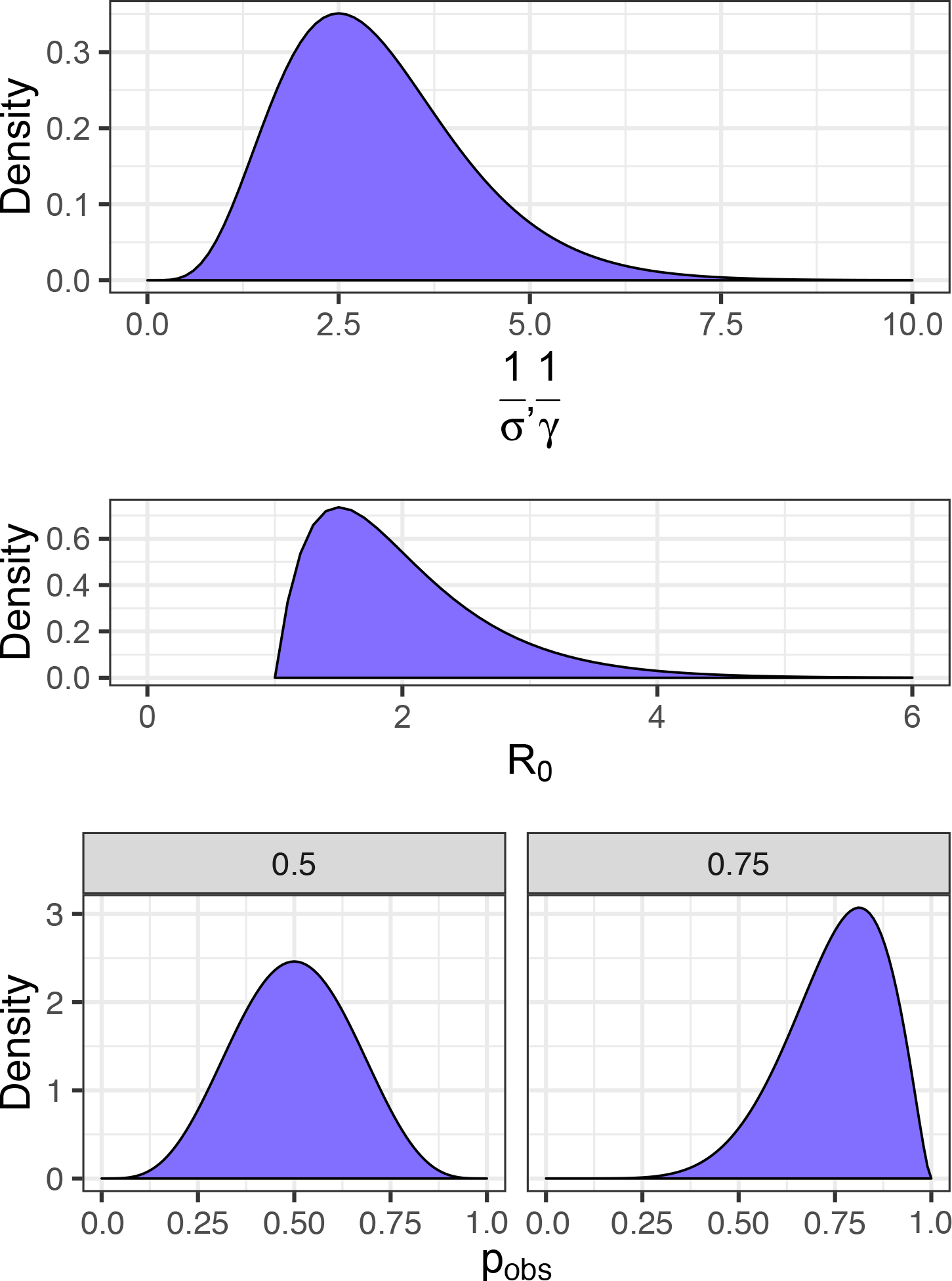
Distributions for model parameters.

**Figure S2:**
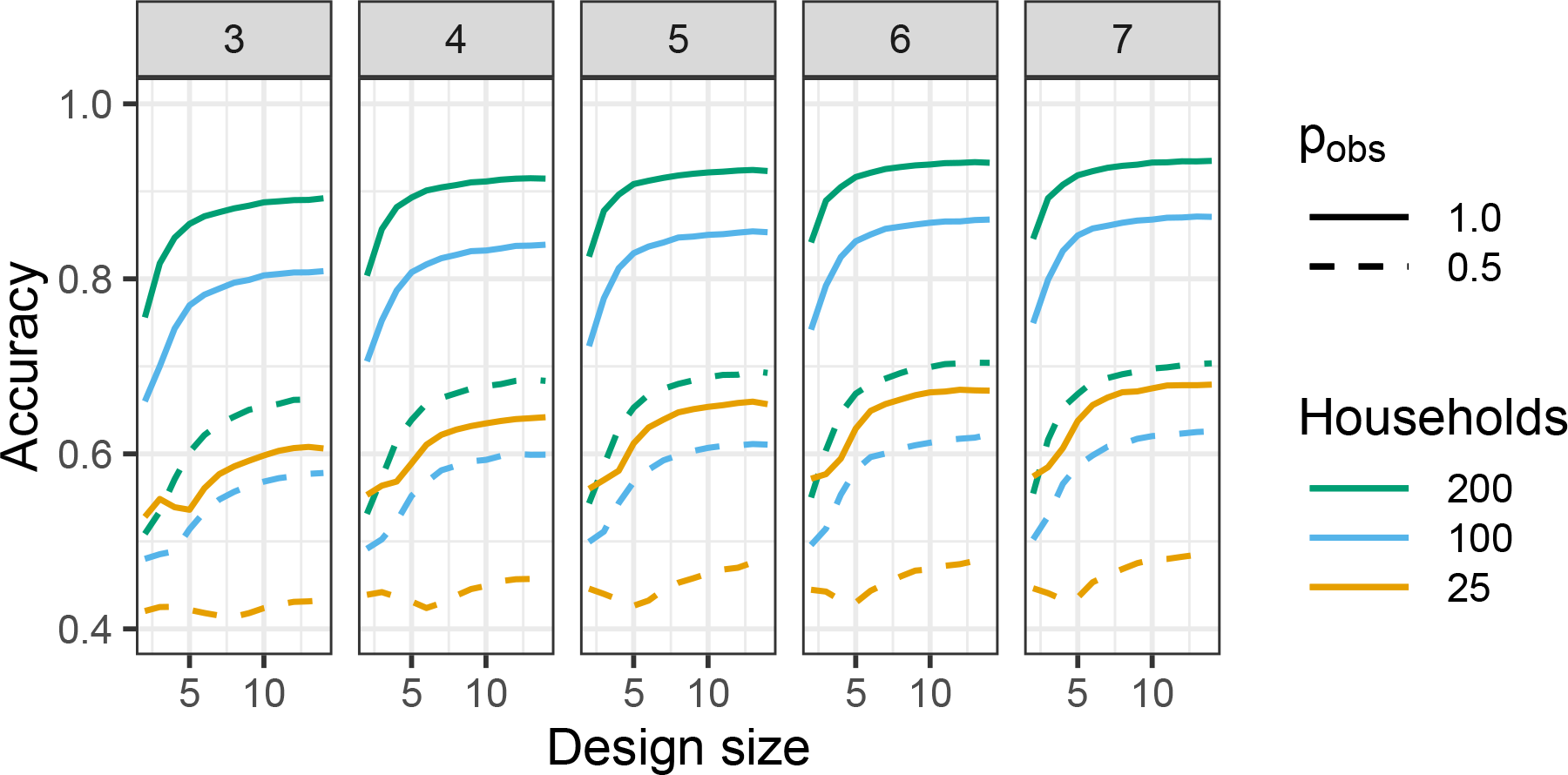
Change in accuracy of optimal designs as household size increases from 3 to 7, under complete observation and mean *p*_obs_ = 0.5. Based on 10,000 training simulations and parameters sampled from distributions.

**Figure S3:**
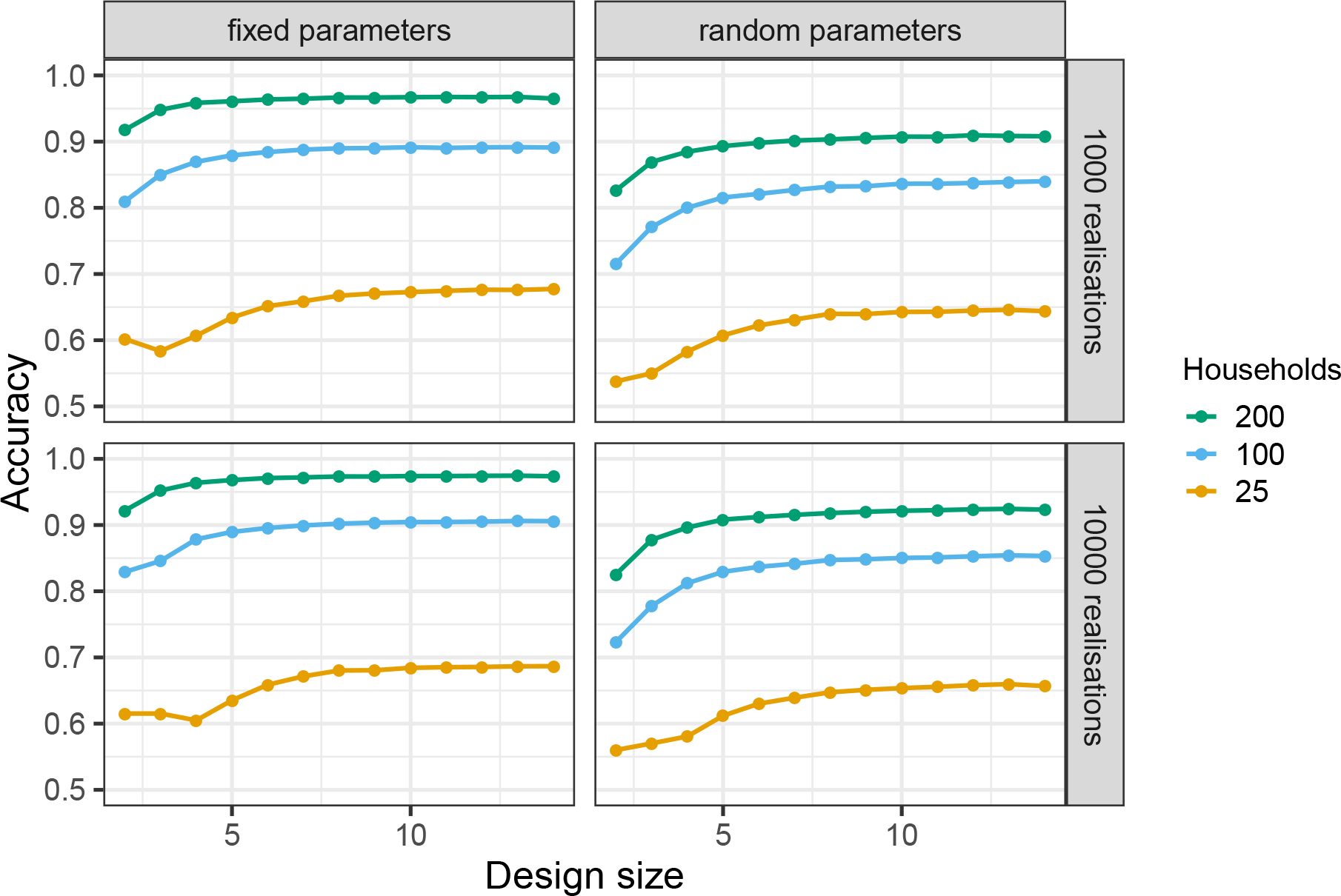
Impact on accuracy of optimal designs as with fixed parameters vs. parameters sampled from distributions, and with 1,000 vs. 10,000 training samples. Based on households of size 5.

**Figure S4:**
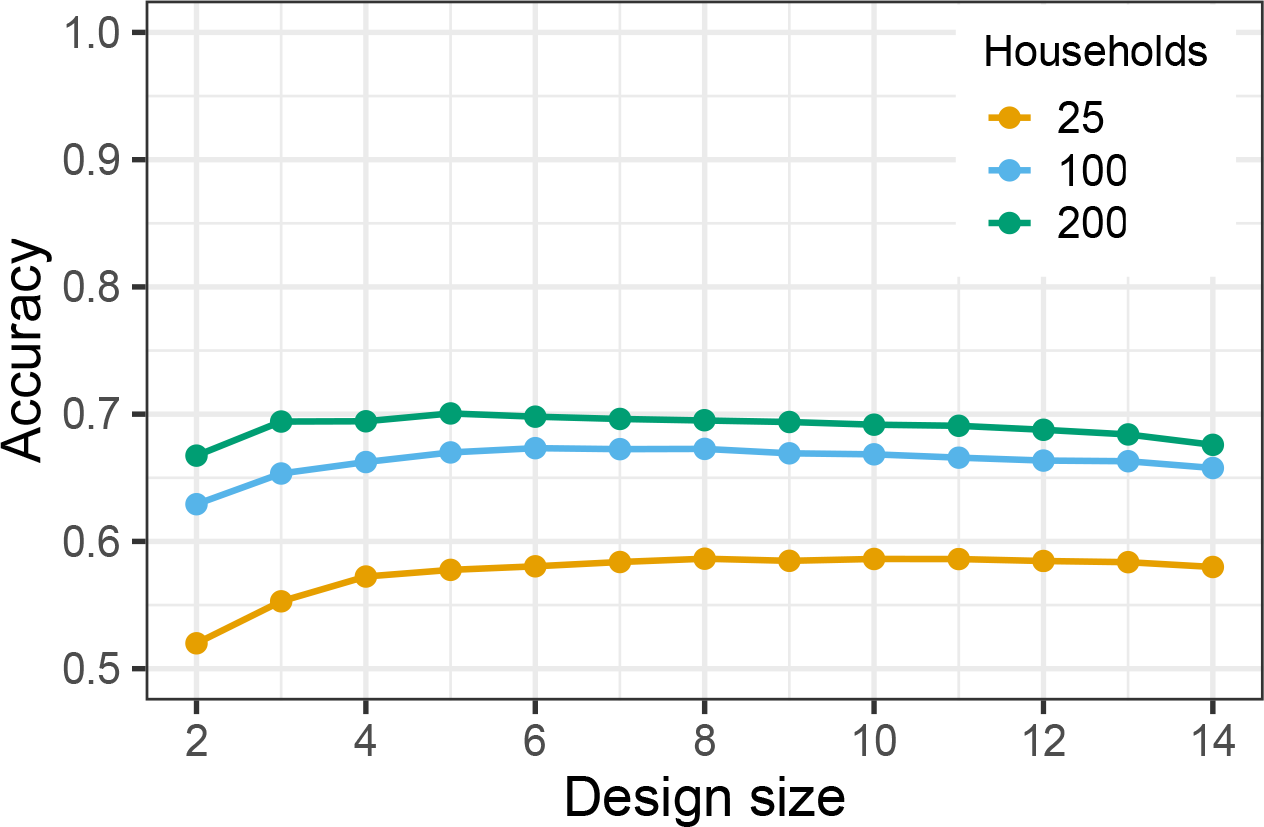
Performance of random forest model discrimination when raw data were used as predictors, rather than histogram summaries (with results in Figure 1d). Based on households of size 5, with 10,000 training points, and random parameters.

## References

1. N. Alzahrani, P. Neal, S.E. Spencer, T.J. McKinley and P. Touloupou Model selection for time series of count data. Computational Statistics Data Analysis 122, 33–44 (2018).

2. R.M. Anderson, C. Fraser, A.C. Ghani, C.A. Donnelly, S. Riley, N.M. Ferguson, G.M. Leung, T.H. Lam and A.J. Hedley Epidemiology, transmission dynamics and control of SARS: the 2002-2003 epidemic. Philos Trans R Soc Lond B Biol Sci. 359, 1091–1105 (2004).

3. Australian Health Management Plan for Pandemic Influenza. http://www.health.gov.au/internet/main/publishing.nsf/content/519F9392797E2DDCCA257D47001B9948/%24File/AHMPPI.pdf (accessed 22/02/19).

4. A.J. Black, N. Geard, J.M. McCaw, J. McVernon and J.V. Ross Characterising pandemic severity and transmissibility from data collected during first few hundred studies. Epidemics 19, 61–73 (2017).

5. A. Black, T. House, M.J. Keeling and J.V. Ross Epidemiological consequences of household-based antiviral prophylaxis for pandemic influenza. Journal of the Royal Society Interface 10, 20121019 (2013).

6. K. Chaloner and I. Verdinelli Bayesian experimental design: a review. Stat. Sci. 10, 273304 (1995).

7. N. Chopin, P.E. Jacob and O. Papaspiliopoulos SMC2: an efficient algorithm for sequential analysis of state space models. Journal of the Royal Statistical Society: Series B (Statistical Methodology) 75, 397–426 (2013).

8. T. Day, A. Park, N. Madras, A. Gumel and J. Wu When Is Quarantine a Useful Control Strategy for Emerging Infectious Diseases?, American Journal of Epidemiology 163, 479485, https://doi.org/10.1093/aje/kwj056 (2006).

9. M.B. Dehideniya, C.C. Drovandi and J.M. McGree Optimal Bayesian design for discriminating between models with intractable likelihoods in epidemiology. Computational Statistics Data Analysis 124, 277–297, https://doi.org/10.1016/j.csda.2018.03.004 (2018).

10. C.C. Drovandi and R.A. McCutchan Alive SMC2: Bayesian model selection for low-count time series models with intractable likelihoods. Biometrics 72, 344–353 (2016).

11. C. Fraser, S. Riley, R.M. Anderson and N.M. Ferguson Factors that make an infectious disease outbreak controllable. Proc. Natl. Acad. Sci. U.S.A. 101(16): 61466151 (2004).

12. A.B. van Gageldonk-Lafeber, M.A. van der Sande, A. Meijer, I.H. Friesema, G.A. Donker, J. Reimerink et al. Utility of the first few100 approach during the 2009 influenza A(H1N1) pandemic in the Netherlands. Antimicrob. Resist. Infect. Control. 1:30 (2012).

13. M. Hainy, D.J. Price, O. Restif and C. Drovandi Optimal Bayesian design for model discrimination via classification. https://arxiv.org/abs/1809.05301 (2018).

14. Y.H. Hsieh, C.C. King, C.W. Chen, M.S. Ho, J.Y. Lee, F.C. Liu, Y.C. Wu, et al. Quarantine for SARS, Taiwan. Emerging infectious diseases 11, 278–82 (2005).

15. M.J. Keeling and J.V. Ross On methods for studying stochastic disease dynamics. Journal of the Royal Society Interface 5, 171–181 (2008).

16. T.G. Ksiazek, E. Erdman, C.S. Goldsmith, S.R. Zaki, T. Peret, S. Emery, S. Tong, C. Urbani, J.A. Comer, W. Lim, et al. A Novel Coronavirus Associated with Severe Acute Respiratory Syndrome. N. Engl. J. Med. 348, 1953–1966 (2003).

17. L.L. Lau, B.J. Cowling, V.J. Fang, K.H. Chan, E.H. Lau, et al. Viral shedding and clinical illness in naturally acquired influenza virus infections. J Infect Dis 201, 15091516 (2010).

18. N. Lee, D. Hui, A. Wu, P. Chan, P. Cameron, G.M. Joynt, A. Ahuja, M.Y. Yung, C.B. Leung, K.F. To, et al. A Major Outbreak of Severe Acute Respiratory Syndrome in Hong Kong. N. Engl. J. Med. 348, 1986–1994 (2003).

19. E. McLean, R.G. Pebody, C. Campbell, M. Champerland et al. Pandemic (H1N1) 2009 influenza in the UK: Clinical and epidemiological findings from the first few hundred (FF100) cases. Epidemiology & Infection 138, 1531–1541 (2010).

20. C. Molnar Interpretable Machine Learning: A guide for making black box models explainable. https://christophm.github.io/interpretable-ml-book/ (2019).

21. E. Patrozou and L.A. Mermel influenza transmission occur from asymptomatic infection or prior to symptom onset? Public Health Rep 124, 193196 (2009).

22. P. Pudlo, J-M. Marin, A. Estoup et al. Reliable ABC model choice via random forests. Bioinformatics 32, 859–866 (2015).

23. F. Pedregosa, G. Varoquaux, A. Gramfort et al. Scikit-learn: Machine Learning in Python. Journal of Machine Learning Research 12, 2825–2830 (2011).

24. L.E. Raileanu, and K. Stoffel. Theoretical comparison between the gini index and information gain criteria. Annals of Mathematics and Artificial Intelligence 41(1), 77–93 (2004).

25. C.P. Robert, J-M. Cornuet, J-M. Marin and N.S. Pillai Lack of confidence in approximate Bayesian computation model choice. Proceedings of the National Academy of Sciences 108, 15112–15117; DOI: 10.1073/pnas.1102900108 (2011).

26. C.P. Robert Approximate Bayesian Computation: A Survey on Recent Results. In: Cools R., Nuyens D. (eds) >Monte Carlo and Quasi-Monte Carlo Methods. Springer Proceedings in Mathematics Statistics, vol 163. (Springer, Cham. 2016).

27. J.V. Ross, T. House and M.J. Keeling Calculation of disease dynamics in a population of households. PLoS ONE 5: e9666 (2010).

28. E.G. Ryan, C.C. Drovandi, J.M. McGree and A.N. Pettitt A review of modern computational algorithms for Bayesian optimal design. Int. Stat. Rev. 84, 128–154 (2015).

29. P. Touloupou, N. Alzahrani, P. Neal, S.E.F. Spencer and T.J. McKinley Efficient model comparison techniques for models requiring large scale data augmentation. Bayesian Anal. 13, 437–459 (2018).

